# Stage-mediated priority effects and season lengths shape long-term competition dynamics

**DOI:** 10.1101/2020.08.14.251926

**Authors:** Heng-Xing Zou, Sebastian J. Schreiber, Volker H.W. Rudolf

**Affiliations:** Program in Ecology and Evolutionary Biology, Department of BioSciences, Rice University, Houston, TX 77005; Department of Evolution and Ecology, University of California, Davis, CA 95616

**Keywords:** priority effects, stage structure, competition, life history, phenology

## Abstract

The relative arrival time of species can affect their interactions and thus determine which species persist in a community. Although this phenomenon, called priority effect, is widespread in natural communities, it is unclear how it depends on the length of growing season. Using a seasonal stage-structured model, we show that differences in stages of interacting species could generate priority effects by altering the strength of stabilizing and equalizing coexistence mechanisms, changing outcomes between exclusion, coexistence, and positive frequency dependence. However, these priority effects are strongest in systems with just one or a few generations per season and diminish in systems where many overlapping generations per season dilute the importance of stage-specific interactions. Our model reveals a novel link between the number of generations in a season and the consequences of priority effects, suggesting that consequences of phenological shifts driven by climate change should depend on specific life histories of organisms.

## Introduction

The relative timing of species’ arrival during community assembly often determines the outcome of species interactions. Generally termed priority effects, this phenomenon is widespread in animal, plant, and microbe systems (Alford and Wilbur 1985; Drake 1991; Louette and De Meester 2007; Kardol et al. 2013; Rasmussen et al. 2014; Clay et al. 2019) and plays a key role in structuring community composition (Chase 2003), species diversity (Fukami 2004), and ecosystem function (Burkle and Belote 2015; Weidlich et al. 2018; Bittleston et al. 2020). Classic and recent studies have highlighted the importance of priority effects in seasonal communities by demonstrating the role of seasonal timing, or phenology, in determining outcomes of species interactions (Sutherland 1974; Morin 1984; Alford and Wilbur 1985; Alexander and Levine 2019; Rudolf 2019; Blackford et al. 2020; Waters et al. 2020). However, despite important advances in identifying mechanisms promoting priority effects (Chase 2003, 2010; Fukami 2004; Tucker and Fukami 2014; Grainger et al. 2019), we still have a poor understanding of how priority effects influence the long-term dynamics and structure of seasonal communities across systems with different life histories. Yet, as climate change shuffles the timing of species arrival (Parmesan 2006 and creating opportunities for priority effects (Rudolf 2019), making this prediction is crucial to fully understand how natural communities respond to a changing climate and which species are most vulnerable.

While various mechanisms have been proposed for priority effects, many are generated by the physiological differences between interacting species that arise by arriving earlier or later. For instance, differences in arrival time alter the traits of species that determine competition, such as resource competition mediated by the size difference between early and late arrivers (Lawler and Morin 1993; Schwinning and Weiner 1998; Brown and Cahill Jr 2022), behavioral interference on late arrivers (Van Buskirk 1992; Poulos and McCormick 2014), altered defense of prey/host of the early-arriving predator/pathogen (Hoverman and Relyea 2008; Hoverman et al. 2013), or even size-dependent intraguild predation (Rasmussen et al. 2014; Sniegula et al. 2019). Here, the outcome of interactions strongly depends on the difference in species’ arrival times (relative arrival times or phenological differences) because timing determines the per-capita effects of interacting species which in turn determines the persistence and coexistence of species. Collectively termed trait-dependent priority effects (Zou and Rudolf 2023), they describe changes in per-capita effects mediated by a shift in the traits of interacting species, and priority effects arise because the shift in timing determines what outcome is possible.

Trait-dependent priority effects represent a category that is different from priority effects defined by classic theory. In traditional models priority effects arise because the per-capita effects of interacting species result in bistability (alternative states) due to positive frequency dependence (species’ growth rate scales positively with its relative abundance; Ke and Letten 2018). In this case, priority effects arise when the more frequent species is always able to maintain a numeric advantage over other species regardless of relative arrival time (phenology). Therefore, the difference in arrival time only matters if it allows the early arriver’s population to increase relative to the late arriver and thus gain a numeric advantage. Traditionally, this type of “frequency-dependent” priority effect has dominated ecological theory on long-term dynamics and species coexistence, assuming no effects of seasonal reassembly on community composition (e.g., Fukami and Nakajima 2011; Gerla and Mooij 2014; Ke and Letten 2018). Although trait-dependent priority effects have been known for decades in empirical systems, they have received surprisingly little attention until recently, as empirical studies find more mechanisms and consequences of trait-dependent priority effects (Rudolf 2018; Blackford et al. 2020; Fragata et al. 2022). Therefore, we are only starting to understand when and how these different types of priority effects influence the long-term dynamics of seasonal communities and species coexistence, especially when phenology varies over seasons and causes variation of community compositions between seasons (Rudolf 2019).

In nature, the importance of trait-dependent priority effects can depend on the length of the growing season (hereafter “season”), specifically on how many generations one season can accomodate (Figure 1). We define a season as the period of growth separated by regular environmental disturbances. If a season consists of only one or a few generations (e.g., annual grassland with a single, non-overlapping generation), trait-dependent priority effects persist because the difference in interacting traits (e.g., ontogenetic stages) is preserved until the end of the season (Figure 1A; Rudolf 2019; Blackford et al. 2020). On the other hand, if many overlapping generations take place in one season (e.g., some zooplankton communities), the correlation between initial arrival time and trait should be weakened because individuals from multiple generations with different trait values interact simultaneously (Figure 1B). In the latter scenario, the long-term outcome of competition should depend less on relative arrival time but more on initial abundances (Toju et al. 2018) and average interaction strengths. Thus, the relative importance of trait-dependent priority effects on the long-term dynamics of communities could vary across systems with different numbers of generations per season.

**Figure 1:**
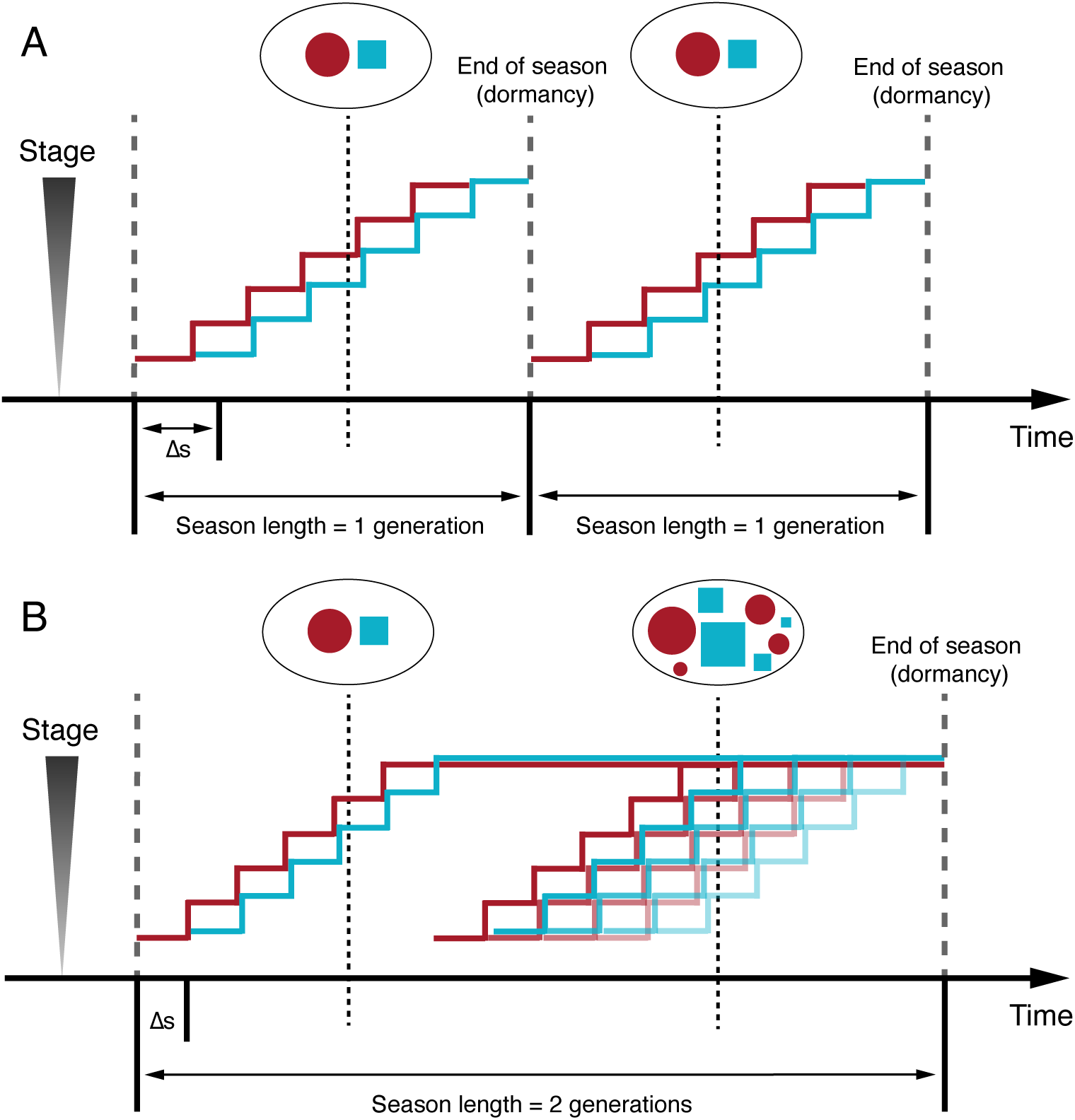
Conceptual drawing of interacting individuals at a given time with single (A) or multiple overlapping (B) generations in a season, indicated by “season length” and thick dashed lines. If a season has only one generation, then the initial size/stage differences caused by different arrival times (phenological difference) persist until the end of the season. If a season is long enough for completion of multiple generations (two shown in B) and individuals continue to reproduce once they mature, populations gradually become mixtures of all different size classes, diluting the effect of initial phenological difference on species interactions. coloured circles and squares indicate two different species and their size/stage distributions. Thick coloured lines show the size/stage of each generation; blue lines in B are slightly raised for visibility. Thin dashed lines shows the community composition at a particular time. In both A and B Δ*s* = *−*1 (species 1 arrives early by one time step); figures not drawn to scale.

To explore the role of trait-dependent priority effects in seasonal systems, we constructed a two-species, stage-structured model to examine how priority effects contribute to the long-term competition dynamics. Specifically, we assumed that the interspecific competition is mediated by stage classes of the two species (“stage-mediated interspecific competition”). We asked how changing relative arrival time would shift outcomes of competition, and how life history (number of generations per season) would affect this shift. We then investigated the contribution of trait-dependent priority effects by including or excluding stage-mediated interspecific competition. To examine the long-term dynamics of seasonal communities in nature, we further investigated the effect of inter-seasonal variation in phenology by allowing arrival times of species to fluctuate across seasons. Together, this allowed us to examine how relative arrival time and season length interact to mediate the importance of priority effects and their role in the long-term dynamics of communities and ultimately persistence and coexistence of competing species.

## Methods

### Stage-structured Model

To capture the potential change in traits of interacting species with different arrival times, we used an age-classified matrix population model with a Beverton-Holt type density dependence (Travis et al. 1980; Caswell 2006). In this model, the age/stage of interacting species is the trait that changes with arrival time; the early-arriving species is at a later stage when the competing species arrives, creating the potential for stage-mediated priority effects (Figure 1; Yang and Rudolf 2010). We included five juvenile stages, one adult stage, and one dormant stage. With a fixed developmental period, using more stages gives a finer scale of arrival time, but does not affect the underlying mechanism of stage-mediated interspecific competition. The dormant stage accounts for individuals before they arrive within a season.

We tracked the seasonal process and species arrival with a function of time: *a_i_*(*t*) = 1 when species *i* is in the community, and *a_i_*(*t*) = 0 when it is dormant. At the beginning of each season, all individuals of species *i* enter the community as stage 1 juveniles with density *N_i,_*_1_. Juveniles in stage *k* of species *i* have density *N_i,k_* and survive to the next stage with probability *P_i,k_*. Adults (*N_i,_*_6_) can live for more than one time step, with a survival probability of *V*, and produce stage 1 juveniles with a density-independent fecundity (*R*). At the end of the season, adults produce offspring that are subject to mortality (*µ*); all other individuals die. These offspring then enters a dormant stage (*N_i,_*_7_) and re-emerge at the assigned arrival time of the next season, with no additional mortality. We let *N_i_* denote the vector of densities (*N_i,_*_1_*, …, N_i,_*_7_) for species *i*.

Overall, the population dynamics of species *i* can be described as:

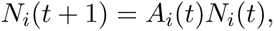

where

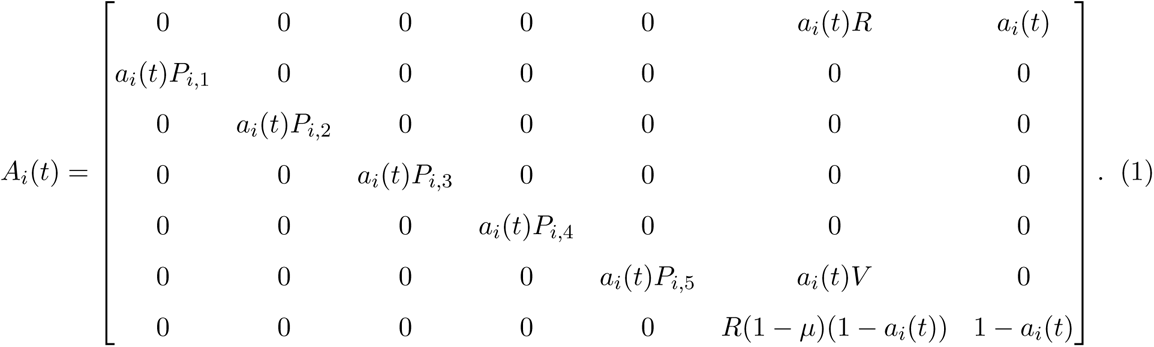

We let Δ*s* denote the difference between the arrival time of species 2 and species 1. Δ*s* = 1 means that stage *m* individuals of species 1 interact with stage *m* + 1 individuals of species 2. Differences in arrival time lead to corresponding differences in what stages interact first; once both species are present, competition is determined by the stage structure of each species. We let *|*Δ*s| ≤* 4 such that the two species can still interact as juveniles even with the latest arrival while they are still in the same generation (i.e., the early species has not reproduced when the late species arrives).

We let both species finish their life cycles within the season. For simplicity, we measure season length using the number of generations (e.g., *T* = 2, 4, 8 generations). We discuss the calculation of season length in Appendix I.

### Stage-mediated Interspecific Competition

We assumed that competition only occurs during the five juvenile stages. This represents a wide range of organisms that experience an ontogenetic niche shift, such as fish, amphibians, and insects (Werner and Gilliam 1984; Miller and Rudolf 2011; Nakazawa 2015). The transition probability of species 1, stage *m* to the next stage is reduced from baseline (*P*_0_) by intra- and interspecific competition from all other individuals:

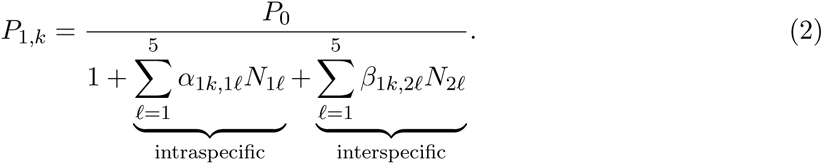

Following a previous approach (Rudolf 2019), the strength of interspecific competition scales according to a sigmoidal function with midpoint (*d*), maximum interspecific competition (*B*) and a scaling constant that determines its shape (*c*):

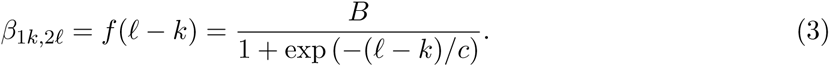

For *β*_2_*_k,_*_1_*_l_* (interspecific competition coefficient representing the effect of stage *l* of species 1 on stage *k* of species 2), the sign of *c* is flipped to create a scaling function of the same shape but in a different direction (Figure 2A). This relationship assumes that later stages are more competitive than early stages, is well supported in plant and animal systems (Shorrocks and Bingley 1994; Rudolf 2018; Blackford et al. 2020), and can arise from various mechanisms such as resource preemption (Schwinning and Weiner 1998; Connolly and Muko 2003) or interference competition (Rudolf 2006; Narvaez et al. 2020). We assumed that the two interspecific competition coefficients are equal when the two species arrive simultaneously. Without this stage-mediated interspecific competition, *β* values are independent of Δ*s*, and all stages should be treated the same; this is equivalent to Δ*s* = 0, leading to *β* = *B/*2. Furthermore, Eqn. 3 is centered at the point (0*, B/*2) and has a range of (0*, B*), meaning that *β* = *B/*2 is the average interspecific competition coefficient if all stages are equally abundant (Figure 2A). Using Eqn. 3 this way ensures the consistency between scenarios with and without stage-mediated interspecific competition. We imposed a competitive hierarchy by setting different intraspecific competition coefficients, i.e., species with lower intraspecific competition is competitively superior (Table 1). We specifically chose intraspecific competition coefficients much smaller than the average interspecific competition coefficient, *B/*2, to promote frequency-dependent priority effects, although higher values did not qualitatively change our results (Appendix III).

**Figure 2:**
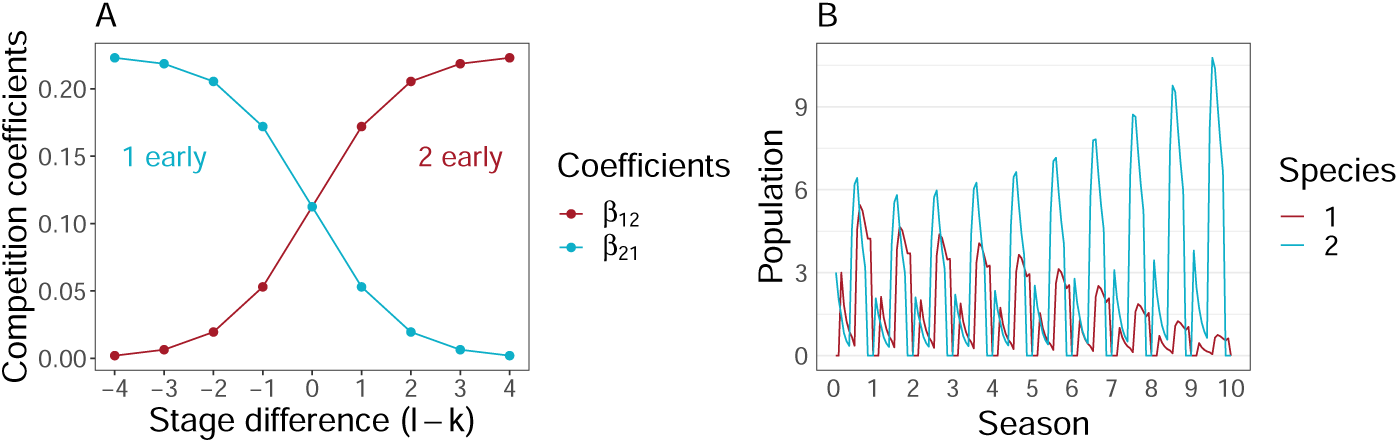
A. The scaling function of stage-mediated interspecific competition. Both per-capita competition coefficients are determined by Δ*s*. The function assumes an early arriver advantage. B. An example of population dynamics generated by the model, with initial density of both species at 3, Δ*s* = 2 (species 2 arrives early by 2 time steps), and *T* = 2 generations. Vertical dashed line marks the beginning of each season. See Table 1 for other parameters used.

**Table 1:** Major parameters and values used in simulation.

| Parameter | Definition | Range/Values |
| --- | --- | --- |
| $\Delta s$ or $\overline{\Delta s}$ | (Mean) initial stage difference | Integers; $[-4, 4]$ (constant)<br>or $[-2, 2]$ (periodic) |
| $v$ | Seasonal variation of $\Delta s$ | 0 (constant) or 2 (periodic) |
| $B, c$ | Baseline interspecific competition, and scaling constant for $f(\Delta s)$, respectively | 0.225, 0, $\pm 0.85$ |
| $\alpha$ | Intraspecific competition coefficient | $\alpha_{11} = 0.05, \alpha_{22} = 0.06$;<br>see results with other values<br>in Appendix III, and results with<br>stage-mediated intraspecific<br>competition in Appendix IV |
| $\beta$ | Interspecific competition coefficient | Determined by $\Delta s$ or<br>$B/2 = 0.1125$ |
| $T$ | Season length | 2, 4, 8, 16, 20 generations |
| $R$ | Fecundity of adults | 12 |
| $S$ | Survival rate of adults after one time step | 0.8 |
| $\mu$ | Seasonal mortality of newly produced $J_1$ | 0.2 |
| $\mathbb{P}_i$ | Baseline survival from juvenile stage $i$ to $i+1$ | 0.8 |

Differences between stages can potentially also alter intraspecific competition and thereby influence the outcome of interspecific competition (Miller and Rudolf 2011; Rudolf and Rasmussen 2013*a*,*b*; Rudolf et al. 2014; Murillo-Rinćon et al. 2017). While including both stage-mediated intra- and interspecific competition is potentially more biologically realistic for a stage-structured system, we present the simpler scenario (constant intraspecific competition coefficients) in the main text for several reasons. Stage-dependent intraspecific competition may be largely irrelevant when conspecific individuals arrive at the same time and have short generation times, but it alters the overall strengths of intraspecific competition when the number of generations per season increases. This change in intraspecific competition can alter the outcome of competition and potential for frequency-dependent priority effects. However, it does not change the qualitative patterns (i.e. interaction of seasonal length and trait-dependent priority effects and we present a detailed analysis of this scenario in Appendix IV). Keeping intraspecific competition coefficients independent of stage structure allows us to isolate the net effect arrival time and season length on interspecific competition. This simplification also greatly facilitates the direct comparison of frequency- vs trait-dependent priority effects scenarios.

### Quantifying Competitive Outcomes

We determined outcomes of competition by calculating the invasion growth rates of each species (MacArthur and Levins 1967; Hofbauer and Sigmund 1998; Schreiber 2000; Roth et al. 2017; Chesson 2018). The invasion growth rate *r_ij_* of species *i* when species *j* is the resident corresponds to its average per-capita growth rate when it is rare and the other species *j*≠*i* is common. We restricted to parameters where each species can persist in the absence of interspecific competition (see Appendix I for details). Through the classic mutual invasiability criterion (MacArthur and Levins 1967; Roth et al. 2017), the invasion growth rates can be used to classify competitive outcomes into three types. When both invasion growth rates are positive (*r*_12_ *>* 0 and *r*_21_ *>* 0), both species increase from being rare and, therefore, coexist. When both invasion growth rates are negative (*r*_12_ *<* 0 and *r*_21_ *<* 0), there is a frequency-dependent priority effect in which the more common species tends to exclude the less common species. When the invasion growth rates are of opposite sign (e.g., *r*_12_ *>* 0 *> r*_21_), one species tends to exclude the other (e.g., species 1 excludes species 2). The proofs and a detailed description of numerically estimating the invasion growth rates *r*_12_ are discussed in Appendix I. Briefly, we numerically determined the resident’s periodic stable state in the absence of the other species, then approximated the invasion growth rate using a linearization of the other species dynamics and the power method from numerical matrix theory (cf. Roth et al. 2017; Benaïm and Schreiber 2019).

While the signs of the invasion growth rates determine *whether* coexistence occurs or not, they do not provide insights into *why* it does or does not occur. To gain insights about why Δ*s* matters for coexistence, we calculated niche difference (ND) and relative fitness difference (RFD) by calculating the sensitivity to competition (Narwani et al. 2013; Godwin et al. 2020). Larger niche differences stabilise species interactions by increasing intraspecific relative to interspecific competition, while smaller fitness differences prevent one species from dominating the other (Chesson 2018). The sensitivity of species 1 is calculated using the long-term growth rate of its monoculture (*r*_1_) and its invasion growth rate at the stable distribution of species 2 (*r*_12_):

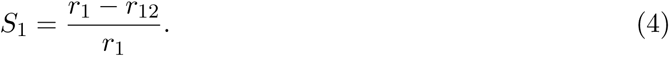

A smaller sensitivity implies higher competitive ability; when both *S <* 1, both species coexist, whereas neither species can invade the other when both *S >* 1. We then used sensitivities of each species to calculate the niche and relative fitness differences (Godwin et al. 2020):

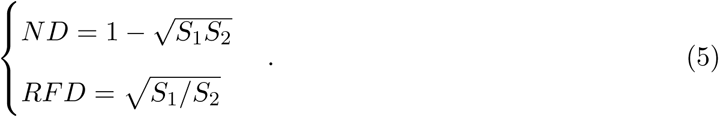

Chesson’s criterion for coexistence requires that:

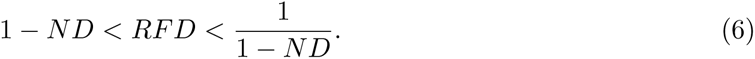

We mapped the calculated ND and RFD values under each Δ*s* and season length onto a “coexistence space” where ranges of coexistence, competitive exclusion, and frequency-dependent priority effects are marked by the above inequality (e.g., Ke and Letten 2018).

### Simulation

All simulations were conducted in R 4.2.1. (R Core Team 2022). We changed season length (*T*), intraspecific competition (*α*), initial stage difference (Δ*s*), and its variation (*v*) to explore their effects on competition outcomes. Each season was at least two-generation long to make sure at least one generation can finish development with a maximum *|*Δ*s|*. We calculated invasion growth rates (*r*) of each species for each parameter combination.

Seasonal variation in arrival time often occurs in natural systems (Sheriff et al. 2011; Diez et al. 2012; Carter et al. 2018) and can have important consequences on long-term community dynamics (Rudolf 2019). We modeled one simple scenario of such variations where species phenology changes periodically over seasons. To achieve this periodicity, we made Δ*s* periodically alternate between 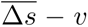 and 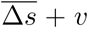 where 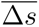 is the mean value and *v* is a positive integer. We required that 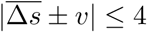, which is the range of Δ*s* explored in simulations without variations. We determined outcomes of competition for using the same method outlined above.

We relaxed several assumptions in our model. First, we allowed the early species to arrive several generations earlier than the late species (Appendix II). Second, we allowed adults to participate in competition and have density-dependent mortality and fecundity (Appendix III). Third, in addition to stage-mediated interspecific competition, we also allowed for intraspecific competition to change with interacting stages (Appendix IV). We report these results in Appendices as they do not qualitatively change our conclusions.

### Code Availability

Full simulation code is deposited at github.com/hengxingzou/Zou2020bioRXiv.

## Results

### Stage-Mediated Priority Effects and Season Length

We found that outcomes of competition were driven by interactions of differences in arrival times and season length. When interspecific competition was stage-mediated, the initial stage difference (Δ*s*) affected the competition outcomes, indicating a stage-mediated priority effect (Figure 3A). When species 1 arrived early (Δ*s <* 0) it generally excluded species 2; when species 2 arrived early (Δ*s >* 0), it either excluded or coexisted with species 1. We expected this asymmetry because we chose ⍺_11_ *<* ⍺_22_ (Table 1), giving species 1 a competitive advantage. However, these stage-mediated priority effects, or changes of competitive outcomes over Δ*s*, were strongly dependent on season length (*T*). When seasons were short (*T* = 2 generations), even the smallest Δ*s* shifted the outcome from alternative stable states (frequency-dependent priority effects) to a single state (competitive exclusion), but this shift in outcome required larger Δ*s* with longer seasons (Figure 3A). As expected, these interactions were contingent on the strength of intraspecific competition. With our model parameters, if both intraspecific competition coefficients were smaller than 0.05, each species generally limited the other more than itself. Longer seasons therefore expanded the region of alternative stable states (Figure S2A). Conversely, ⍺ values larger than 0.06 promoted coexistence, and the two species coexisted regardless of their initial stage differences when each season contained more than 8 generations (Figure S2B). Although we fixed the baseline value of interspecific competiton (*B*) and changed the range of intraspecific competition (⍺) in our model, shifting *B* can have similar effects if its relationship with *α* remains the same (i.e., if a species limits itself more than the other, the set of *α* and *B* will promote coexistence). While specific patterns (e.g., the amount of Δ*s* and the season length at which competitive outcomes are shifted) are contingent on the exact values of intraspecific competition coefficients (Table 1), the general interaction between Δ*s* and season length was always present. We observed that the early arriver sometimes could not exclude the late arriver when the latter arrived extremely late (*|*Δ*s|* = 3, 4) in several scenarios (Figure 3A; Figure S2A). This is because larger the Δ*s*, the lesser the temporal overlap between the two species, and the overall interspecific competition became smaller. If we allowed for the two species to arrive over the time span of more than one generation (*|*Δ*s| >* 4), sufficiently large Δ*s* would eventually lead to coexistence because both species would be increasingly asynchronous. Indeed, increasing Δ*s* strongly increased niche difference between the two species, promoting coexistence (see below). We discussed this late arriver advantage in detail in Appendix II.

**Figure 3:**
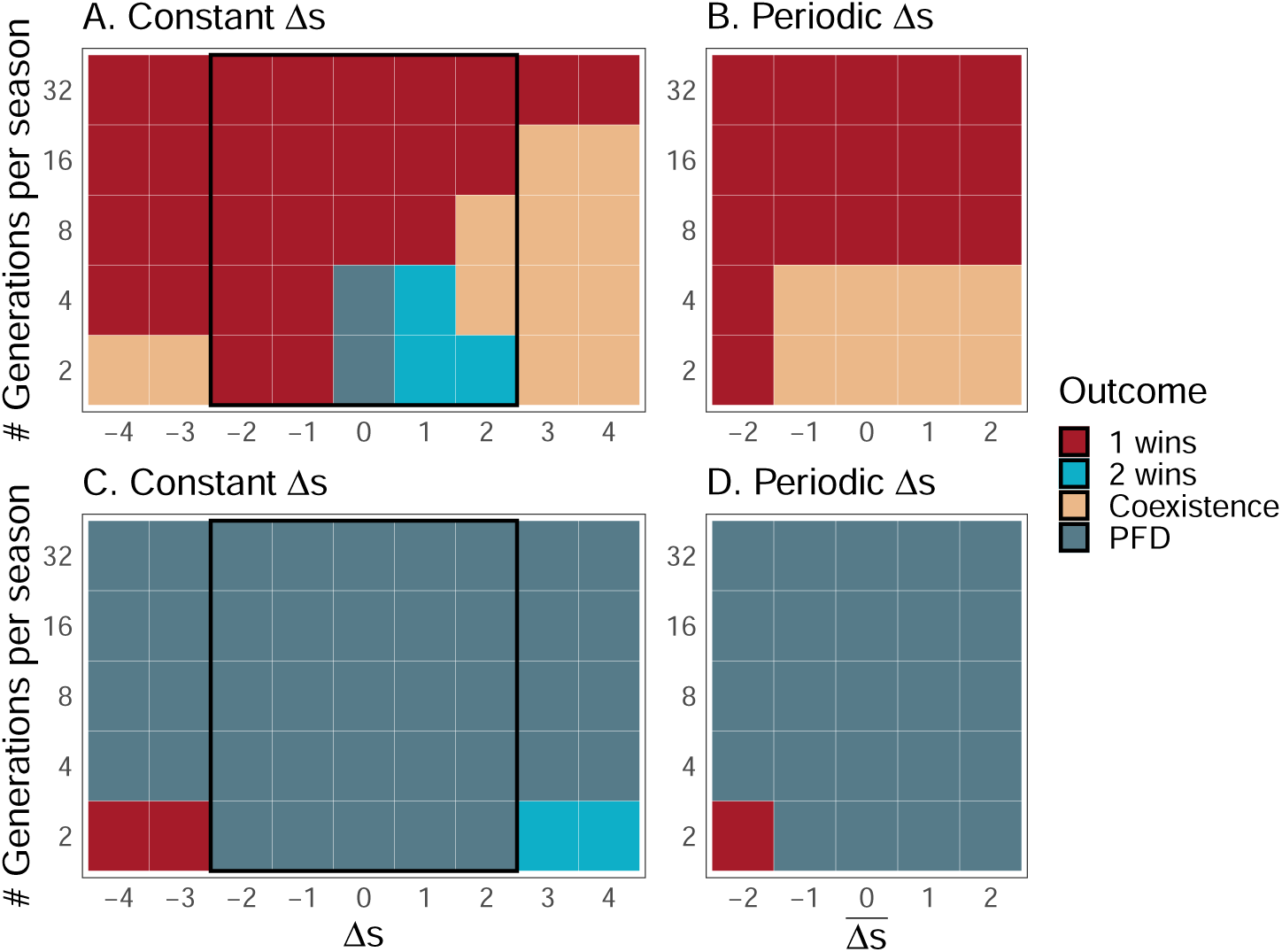
Phase diagrams of competition outcomes across different season lengths (*T* ; in number of generations per seasons) and stage differences (Δ*s*). A and B show results with stage-mediated interspecific competition, C and D show results without stage-mediated interspecific competition. Left panels show results from simulations with constant Δ*s* across seasons. Black rectangles in panels A and C show the parameter space examined by periodic simulations presented in panels B and D. PFD stands for positive frequency dependence (frequency-dependent priority effects). See Table 1 for other parameters used.

When we removed stage-mediated interspecific competition, outcomes were mostly driven by frequency-dependent priority effects (the species with a larger initial population won; Figure 3C), except when *T* = 2 generations, where the early arriver always won when it arrived sufficiently early (Δ*s* = *−*3*, −*4, i.e., early species at stage 4 or 5 when the late species arrived). In these cases, the early arriver was exempted from interspecific competition, and this exemption gave the early arriver an advantage large enough to override the tendency towards positive frequency dependence. In longer seasons, the early arriver advantage became weaker. Overall, the interaction of arrival time and season length indicates that longer seasons remove stage-mediated priority effects. Changing intraspecific competition coefficients did not qualitatively affect this observation (Figure S2).

### Coexistence Space

Phenological shifts (initial stage differences; Δ*s*) and season length (*T*) changed both niche and relative fitness differences between species, further demonstrating the complex nature underlying the early arriver advantage observed in Figure 3. With shorter seasons (*T* = 2 generations), changing the initial stage difference from *−*4 (species 1 at stage 5 when species 2 arrived) to 4 (species 2 at stage 5 when species 1 arrived) strongly shifted niche differences (ND) and relative fitness differences (RFD) from promoting coexistence to species 1 winning, frequency-dependent priority effects, species 2 winning, and coexistence again (Figure 4A). In general, smaller initial stage differences (*|*Δ*s|*) led to a smaller niche difference, meaning that the early arriver had less competitive advantage; when both species arrived together (Δ*s* = 0), the niche difference become negative, indicating a a strong tendency towards positive frequency dependence (Ke and Letten 2018). This is consistent with frequency-dependent priority effect we observed from our simulations. On the other hand, fitness differences between two species generally increased with larger initial stage differences. When the initial stage difference was large (*|*Δ*s|* = 3, 4), the late species was slightly advantageous and was not excluded because of stage-mediated interspecific competition (see above), and this advantage is reflected by fitness differences that were smaller than niche differences, leading to coexistence (Figure 4A). With longer seasons, larger *|*Δ*s|* monotonically increased both niche and fitness differences due to the early arriver advantage (Figure 4B).

**Figure 4:**
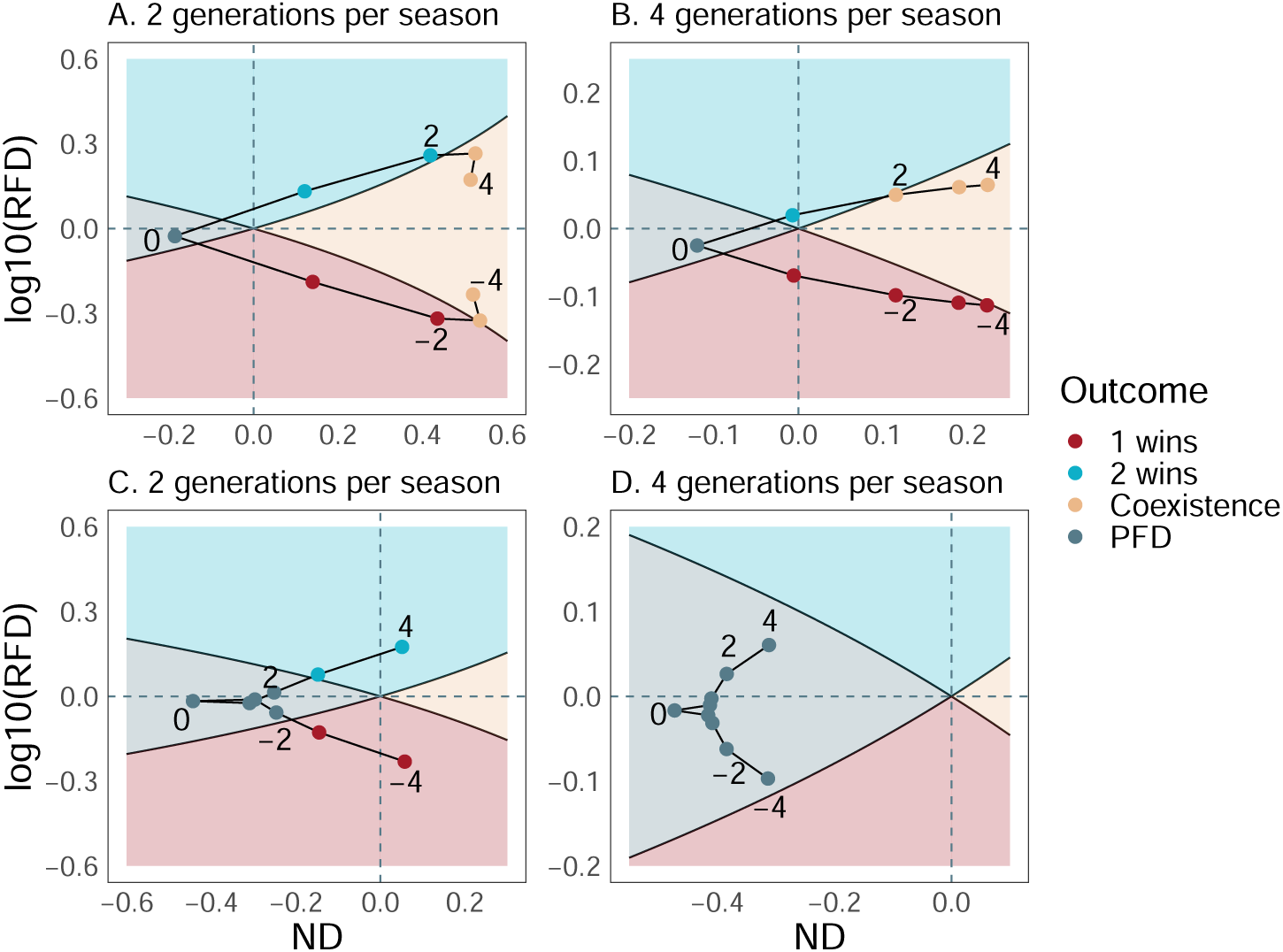
Calculated niche difference (ND) and relative fitness difference (RFD) for interactions at different Δ*s*. Numbers next to points denote values of Δ*s* as it changes from −4 to 4 following the trajectory, showing the movement of ND-RFD points in the coexistence space. colours of each point and shades of the coexistence space indicate regions of each outcome. PFD stands for positive frequency dependence (frequency-dependent priority effects). A and B show results with stage-mediated interspecific competition, C and D show results without stage-mediated interspecific competition. Left panels show results of *T* = 2 generations, right panels show results of *T* = 4 generations. Note that axis ranges of each panel are different for better visibility, but corresponding panels in Figure 4 and 5 have the same axes for better comparison between results without and with periodic fluctuations. See Table 1 for other parameters used.

Without stage-structured interspecific competition, we still observed changes in niche and fitness differences with shifts in Δ*s*. Overall, larger initial stage differences increased both niche and fitness differences. With shorter seasons (*T* = 2 generations), shifts in niche and fitness differences were large, and these shifts led to competitive exclusion when *|*Δ*s|* was large (Figure 4C). With longer seasons (*T* = 4 generations), the effect of Δ*s* on the adult population was smaller, and all calculated values fell in the positive frequency dependence region, indicating a strong tendency towards frequency-dependent priority effects (Figure 4D).

Overall, these patterns demonstrated that initial stage differences strongly affected outcomes of competition by changing both niche and relative fitness differences of interacting species. This effect was stronger when seasons were short and became weaker with longer seasons, which allowed for overlapping generations and many coexisting stages.

### Periodic Environment

Periodic variations in initial stage differences (Δ*s*) allowed for each species to arrive early at alternate seasons, shifting outcomes of competition compared to simulations with constant Δ*s*. In most cases with a change of outcomes, these variations switched competitive exclusion or positive frequency dependence to coexistence, although in some cases periodic variations also switched coexistence or positive frequency dependence to competitive exclusion (Figure 3A, 3B). Periodic variations generally led to smaller ranges of niche and fitness differences. With stage-mediated interspecific competition, niche differences decreased when the average arrival times of species were most different (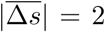; Figure 5); this equalizing force arose because the species alternated in being the early arriver. However, niche differences increased when the two species on average arrive at the same time 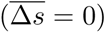 because even though the average arrival times remained the same, the periodic variations increased the chance that one species arrived early. With stage-mediated interspecific competition, periodic Δ*s* decreased the relative fitness difference. Without stage-mediated interspecific competition, periodic variations of Δ*s* slightly decreased relative fitness differences and made niche differences slightly less negative, but they mostly promoted frequency-dependent priority effects, except when species 1 arrived much earlier at the shortest season (*T* = 2 generations; Figure 3D), which was caused by the slight increase in the relative fitness difference (Figure 5C). Overall, These shifts in equalizing and stabilizing mechanisms corresponded to observed changes in long-term competitive outcomes.

**Figure 5:**
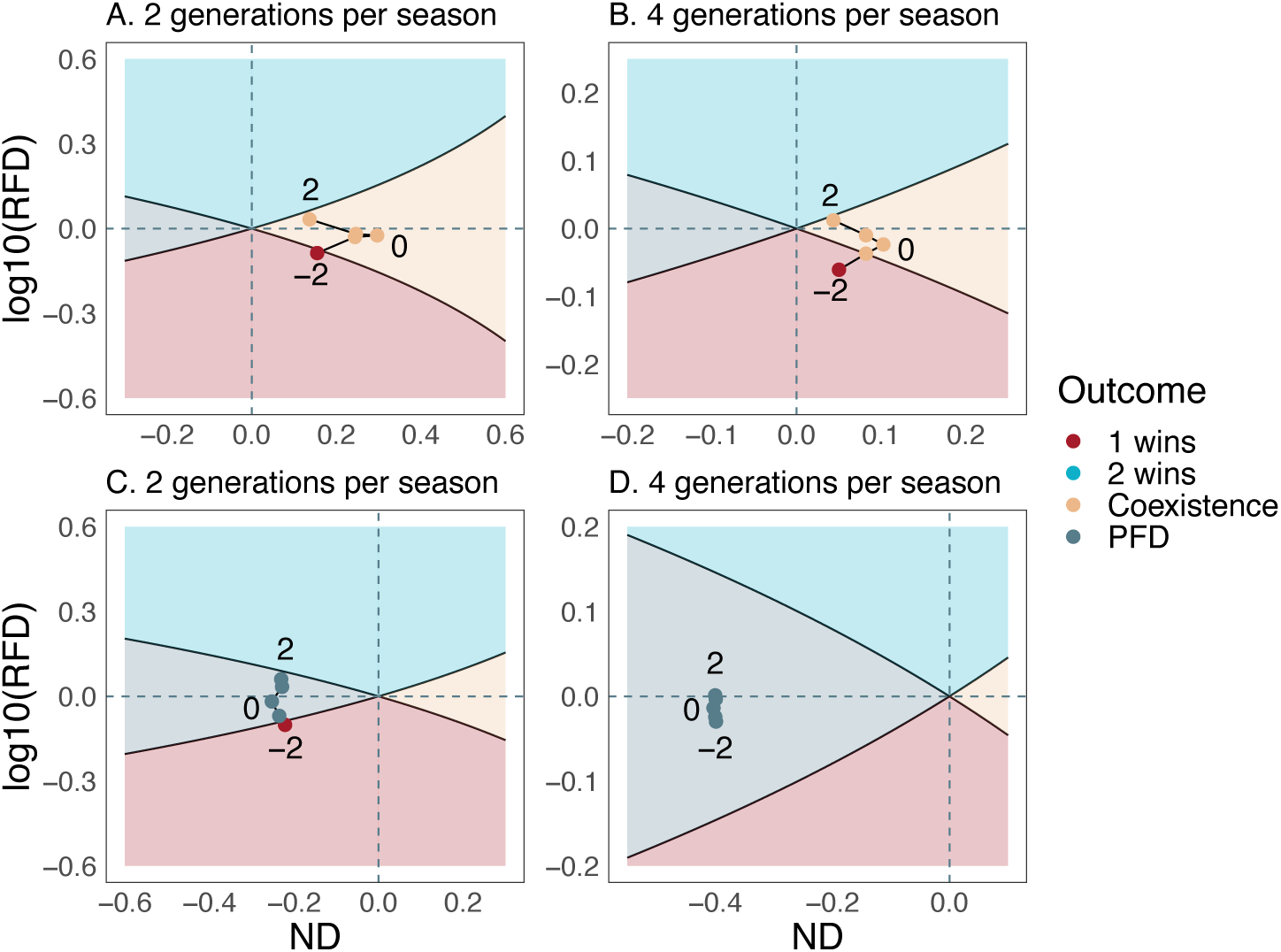
Calculated niche difference (ND) and relative fitness difference (RFD) for interactions at different Δ*s* under periodic fluctuation. Numbers next to points denote values of Δ*s* as it changes from −2 to 2 following the trajectory, showing the movement of ND-RFD points in the coexistence space. colours of each point and shades of the coexistence space indicate regions of each outcome. PFD stands for positive frequency dependence (frequency-dependent priority effects). A and B show results with stage-mediated interspecific competition, C and D show results without stagemediated interspecific competition. Left panels show results of *T* = 2 generations, right panels show results of *T* = 4 generations. Note that axis ranges of each panel are different for better visibility, but corresponding panels in Figure 4 and 5 have the same axes for better comparison between results without and with periodic fluctuations. See Table 1 for other parameters used.

## Discussion

Priority effects can greatly influence community structure and functions (Chase 2003; Fukami 2004; Burkle and Belote 2015; Weidlich et al. 2018; Bittleston et al. 2020), but their long-term implications on seasonal community dynamics and underlying mechanisms are poorly understood. Using a stage-structured competition model that accounts for the seasonal nature of ecosystems, we show that the phenological differences within seasons (initial stage difference) and their fluctuations between seasons determine the long-term dynamics of two competing species, generating stage-mediated priority effects. Furthermore, these effects are dependent on the life histories of competing species (number of generations per season). In systems with only one or a few generations per season, changes in mean and seasonal variation of relative arrival time (phenology) at the onset of the season led to stage-mediated priority effects that altered the outcome of interactions. This effect was dampened and even disappeared in systems with many overlapping generations per season. Thus, systems with less generations per season are much more sensitive to climate-mediated changes in phenology. Together, these results suggest that different mechanisms of priority effects could be important for systems operating at different time scales, and emphasise the role of seasonality, phenological shifts, and variations in long-term community dynamics.

### Effects of Relative Arrival Time and Season Length

Phenological differences could lead to stage-mediated priority effects in various systems (Lawler and Morin 1993; Geange and Stier 2010; Rasmussen et al. 2014; Alexander and Levine 2019; Blackford et al. 2020; Fragata et al. 2022), but the long-term consequences of these effects remain poorly resolved. Recent theory suggests that stage-mediated priority effects can play a key role in determining long-term dynamics in seasonal communities, but this work largely ignores species life history such as stage structure and season lengths (Rudolf 2019). By incorporating these biological realisms, we show that long-term consequences of stage-mediated priority effects are strongly influenced by the life histories of species, specifically the number of generations in a season. Indeed, shifting initial stage differences (Δ*s*) in our model changes the niche difference and relative fitness difference between the two species, but this change weakens with more generations per season. By accommodating multiple overlapping generations, longer seasons allow all stages to interact with each other (Figure 1). As the stage distribution of two species become more similar, the initial stage difference becomes less important. Thus, increasing initial stage differences “pushes” the system to a new potential outcome, while increasing season length “pulls” the system back, constraining the range of potential outcomes by reducing the effective trait differences.

Temporal variation can play an important role in shaping the outcome of species interactions (Chesson 2000; Adler et al. 2006; Barabás et al. 2018; Rudolf 2019). In nature, relative arrival time fluctuates across seasons, and this fluctuation of phenology could be increasing with climate change (Parmesan 2006; Post 2013; Pearse et al. 2017; but see Stemkovski et al. 2022). Our results indicate that even the simplest, non-random seasonal fluctuation in relative arrival time could change competition outcomes, especially when stage-mediated mechanisms are present: periodic arrival times enable either species to harness early arriver competition advantage in alternating seasons, contributing to both equalizing and stabilizing mechanisms in the presence of stage-mediated interspecific competition. As in the cases without fluctuations, these effects are strongest with fewer generations per season because of more accentuated stage-mediated priority effects. Thus, how seasonal variations in species’ phenology change long-term community composition is highly dependent on the type of priority effect and species life histories.

Together, these results also suggest that species can systematically differ in the sensitivity to climate-mediated shifts in phenology. Systems with more generations per season should be more buffered against seasonal variation in phenology, as the contribution of relative arrival time to community dynamics decreases with the number of generations per season. In nature, this comparison of less vs. more generations per season may arise from two scenarios. First, individuals of the same species may increase their development rates and voltinism at different climatic conditions, at different geographic regions (e.g., city vs. rural areas; Brans and De Meester 2018) or with long-term climate change (Altermatt 2010; Iler et al. 2021). However, these changes are often on the scale of one or two generations, and may be less likely to change the long-term consequences of species interactions, depending on the demographic rates of the system. Second, in the long term, interacting species within taxa that reproduce multiple times in a season (e.g., zooplanktons, insects) may be less subject to priority effects arising from phenological differences than those in taxa with less generations in a season (e.g., annual plants, amphibians). Previous studies have discovered several mechanisms that could buffer against mismatch arising from phenological variation, such as decreased competition (Reed et al. 2013), intrinsic flexibility in phenology and resource use (Salido et al. 2012), or dispersal across heterogeneous patches (Phillimore et al. 2012; Post 2013). As warming affects life history traits, we need more empirical work to understand the synergistic effects of life histories and phenological shifts.

### Mechanisms of Priority Effects

Until recently, the distinction between frequency- and trait-dependent mechanisms of priority effects has received little attention. The niche-based mechanistic framework of priority effects, namely niche preemption and niche modification (Vannette and Fukami 2014; Fukami 2015), aligns well with our definition of trait-dependent priority effects because changes in interacting traits would likely alter realised niches of both species. Indeed, the calculated niche and relative fitness differences values changed with initial stage differences in our model, demonstrating the effect of relative arrival times on niches. On the other hand, a recently developed quantitative framework of priority effects (Fukami et al. 2016; Ke and Letten 2018) requires fixed competition coefficients regardless of arrival time. It is, therefore, unable to fully address niche-based mechanisms and can only be applied to frequency-dependent priority effects arising from positive frequency dependence. By mapping competition at different initial stage differences onto the coexistence space, our results indicated the difference between frequency- and trait-dependent priority effects. While a species pair exhibiting the former falls into a region in the coexistence space, the latter is marked by a *shift* of niche and fitness differences in this space: greater differences in arrival times generally increase niche difference and absolute values of relative fitness difference, leading to a shift in outcomes of competition. This difference between frequency- and trait-dependent priority effects is highlighted by a recent empirical study that did not find priority effects via positive frequency dependence but via preemption of spatial niches, detected through the shift from competitive exclusion to coexistence (Fragata et al. 2022).

In our model, trait-dependent priority effects are achieved by stage-mediated interspecific competition: interactions are affected by the ontogenetic stages of the two species (Yang and Rudolf 2010; Rudolf 2019). This effect amplifies the early arriver advantage: while arriving early already exempts one species from interspecific competition for several time steps, stage-mediated interspecific competition also allows the early arriver to have a stronger impact on the late arriver. However, we found that trait-dependent priority effects can attenuate and even disappear when longer seasons allow for multiple overlapping generations. This indicates that priority effects may be generated by fundamentally different mechanisms in systems with different numbers of generations per growing season. In systems with only a few generations per season (e.g. annual grassland, amphibians and dragonflies in ephemeral ponds), trait differences caused by relative arrival time may persist and mediate species interactions for the whole season (Rasmussen et al. 2014; Blackford et al. 2020). In systems with many generations per season, priority effects observed are likely generated by positive frequency dependence (Fukami 2004; Grainger et al. 2019); they are less likely to display trait differences (e.g., stage classes) and more likely to reach equilibrium within one season.

The two underlying mechanisms of priority effects could lead to different community dynamics in the long term; one such example is the persistence of priority effects in spatial communities over several dispersal or colonization events. Dispersal could homogenise local community composition regardless of different assembly history, lowering the impact of priority effects (Mouquet and Loreau 2002; Fukami 2015; Grainger and Gilbert 2016; but see Vannette and Fukami 2017). However, theory and experiments found that when trait-dependent priority effects are present, dispersal may maintain community dissimilarity among patches (Pu and Jiang 2015; Miller and Allesina 2021; Zou and Rudolf 2023). These results highlight the need of considering specific mechanisms of priority effects.

### Next Steps

Trait-dependent priority effects are common and can be driven by various mechanisms. Our model considered the ecological differences between ontogenetic stages of interacting species (stage-mediated priority effects), which correlate with relative arrival times and determine species interactions in both animal and plant systems (Alford and Wilbur 1985; Van Buskirk 1992; Lawler and Morin 1993; Rasmussen et al. 2014; Godoy and Levine 2014; Carter and Rudolf 2019; Brown and Cahill Jr 2022). How age or stage is related to changes in interaction strength depends on the specific mechanisms and thus can vary across systems. For instance, the competitive ability may change quickly or slowly with differences in stages in systems with more vs. less generations per season, respectively. While these relationships will determine the quantitative results (e.g., the absolute difference in arrival time needed to change competition outcome, or the number of generations needed to remove the effect of arrival time), they are unlikely to change the general patterns presented here.

Our model is based on several important assumptions. We assumed no competition between adults and no reproduction from juveniles. Competition between adults is relevant to systems with less ontogenetic niche shifts such as plants and zooplankton. We also assumed no stage-mediated competition between conspecific individuals, which may not apply to certain systems. Although relaxing the above two assumptions did not qualitatively change the outcomes in our model (Appendix III, IV), different results may arise using other model formations or in experimental systems. In nature, relaxing either assumption may increase the total strength of competition experienced by smaller-stage or younger individuals from larger stages of both species, and this additional competition may decrease juvenile survival, rate of development, or overall fecundity of the population. These effects will likely influence the initial population of the next season, which may subsequently affect competition outcomes if they are dependent on relative frequencies of the competitors (Zou and Rudolf 2022). Including reproduction of late juvenile stages, such as in some zooplanktons, may promote overlapping generations and possibly further decrease the importance of initial phenological difference. More importantly, we assumed the same number of stages and duration of each stage and therefore similar developmental rates for both species. If larger stages still have a competitive advantage, then stage-mediated priority effects are likely still present. However, species with a faster physiological development may gain an additional advantage. This is especially relevant because climate change may also accelerate vital processes such as ontogenetic development and reproduction timing (Brans and De Meester 2018; Keller and Shea 2020), but how the two contrary effects interact in competitive communities is less explored (Lancaster et al. 2017). Finally, we do not consider the role of rapid evolution. Theory and experiments have found that such rapid adaptation of early arrivers (monopolization effects) can alter long-term competition dynamics by reducing competitive dominance (Leibold et al. 2019; Nadeau et al. 2021). This is especially relevant for systems with many generations per season because species may be able to quickly adapt within a season.

Stage differences are also not the only mechanism that could cause trait-dependent priority effects. For instance, priority effects in plants could arise from light competition (Schwinning and Weiner 1998), plant-soil feedbacks (Kardol et al. 2007; Van der Putten et al. 2013) or plant-pollinator interactions (Waters et al. 2020); arrival time of competing predators or parasites can also alter defensive traits of common prey or host, subsequently affecting competition (Olito and Fukami 2009; Clay et al. 2019). All of these alternative mechanisms could lead to a different competition-phenology function (different shape or direction of curves in Figure 2A) that often is poorly documented or unknown in natural systems. Our study provides a baseline of how trait-dependent priority effects from one specific mechanism (stage-mediated interspecific competition) could affect community dynamics that can easily be expanded to explore the long-term effects of these different systems once more information is available.

Recent attention to phenological shifts (Rudolf 2019) and seasonality (White and Hastings 2020) raises the need of rethinking priority effects in seasonal ecosystems. By incorporating stage-mediated interspecific competition, our model shows that the importance of trait-dependent priority effects depends on the reproduction frequency of the community. These results provide a much-needed link between priority effects and species life histories. As we face pressing global change, we need a better resolution on how communities respond to the simultaneous shifts in phenology and life history. Future studies should continue to investigate such synergistic effects under season- and time-explicit frameworks.

## Acknowledgments

We thank Joshua Fowler, Tom E. X. Miller and Zoey Neale for feedback on the manuscript. We thank Jennifer Lau, David Vasseur, Lin Jiang and six anonymous reviewers for helpful comments on the manuscript. Funding was provided to V.H.W.R by NSF DEB-1655626 and to S.J.S by NSF DEB-2243076.

## Notes

https://github.com/hengxingzou/Zou2020bioRXiv

### Competing Interest Statement

The authors have declared no competing interest.

### Summary of Updates

This version contains updated formation of the simulation model, subsequent results, and changes of terminologies throughout the manuscript.

