## Supplementary Material for "Stage-mediated priority effects and season lengths shape long-term competition dynamics"

### Appendix I. Supplemental Methods

#### Calculating Stable Stage Distribution

We determined the stable stage distribution of the model by numeric simulations, which require that the new population is close or identical to the population of previous time step. Mathematically, this means that the Euclidean distance between population vectors at two time steps is close to 0:

$$d(N_i(t-1) - N_i(t)) < \epsilon \text{ for sufficiently small } \epsilon > 0. \tag{1}$$

We used $\epsilon = 10^{-12}$ as the threshold in simulations. In simulation, we ran the model for indefinite time and calculated the Euclidean distance between the initial population of the latest two seasons, until this distance is smaller than $10^{-12}$.

#### Setting the Length of Each Season

At the end of the assigned number of generations, all species reproduce, and the resulting population all switch to a "dormant" stage until its assigned arrival time of the next season. Therefore, although our stage-structured model consists of seven stages of each species, including the dormant stage, this stage only appears at the end of each season. Season lengths are thus not multiples of seven time steps (the number of stages in the model) but multiples of six, plus an additional time step at the end of each season that accounts for the shift to dormancy, if the two species arrive simultaneously ($\Delta s = 0$). If $\Delta s \neq 0$, each season is extended to let the late arriver finish its final generation. Therefore, the length of each season in simulations is $n * 6 + |\Delta s|$. When $\Delta s = 0$ this translates to $13, 25, 49, 97$, and $193$ time steps, corresponding to $T = 2, 4, 8, 16$, and $32$ generations per season.

#### Numeric Methods for Calculating Invasion Growth Rates

Consider a two species, stage structured, periodic matrix model of the form

$$N_i(t+1) = A_i(t, N_1(t), N_2(t))N_i(t) \tag{2}$$

where $N_i(t)$ are the vectors of population densities, $A_i(t, N_1, N_2) = A_i(t+T, N_1, N_2)$ for all population vectors $N_1, N_2$ and times $t$, i.e., the model is period $T$.

While we are interested in classifying the outcome of competition for both species, we first need to understand whether species 1 (similarly for 2) by itself can persist in the absence of the other. This requires analyzing the submodel

$$N_1(t+1) = A_1(t, N_1(t), \mathbf{0})N_1(t) \tag{3}$$

where the densities of species 2 are set to zero, i.e., $\mathbf{0} = \begin{pmatrix} 0 \\ 0 \\ \vdots \\ 0 \end{pmatrix}$. Persistence is determined by the linearization of the dynamics at $N_1 = \mathbf{0}$ i.e. the density-independent model

$$N_1(t+1) = A_1(t, \mathbf{0}, \mathbf{0})N_1(t) \tag{4}$$

Let $\lambda_1$ be the dominant eigenvalue of the matrix product $\prod_{t=1}^{T} A_1(t, \mathbf{0}, \mathbf{0})$. Then the per-capita growth rate of species 1 when both species are rare is $r_1 = \frac{1}{T}\log\lambda_1$. When $r_1 > 0$, the population persists (in the sense of permanence). When $r_1 < 0$, the extinction equilibrium $N_1 = \mathbf{0}$ is linearly stable and initial conditions near this equilibrium will tend exponentially quickly to extinction. We can define $r_2$ similarly.

For the remainder of this section, we assume $r_1 > 0$ and $r_2 > 0$, i.e., the two species can persist in the absence of interacting with one another. Furthermore, we assume that the population dynamics for the single species always converge to a periodic attractor. Namely, for species 1, there is a period $T$ solution, $N_1^*(0), N_1^*(1), N_1^*(2), \ldots$ with $N_1^*(t) = N_1^*(t+T)$ for all $t \geq 0$, and for any other solution $N_1(t)$ with $\sum_j N_{1j}(0) > 0$, there is a phase $0 \leq \tau \leq T-1$ (depending on the initial condition) such that

$$\lim_{t\to\infty} \|N_1(t+\tau) - N_1^*(t)\| = 0$$

Similarly, we assume there is globally stable period $T$ solution $N_2^*(t)$ for single species 2 model.

To classify the ecological dynamics of the two species model, we consider the cases where one species is (infinitesimally) rare and the other is more common. The first species is called the invader and the other, the resident. Let us assume species 1 is the resident and species 2 is the invader. While the invader remains rare we can approximate the community dynamics with the partially linearized system

$$\begin{aligned} N_1(t+1) =& A_1(t, N_1(t), \mathbf{0})N_1(t) \\ N_2(t+1) =& A_2(t, N_1(t), \mathbf{0})N_2(t) \end{aligned} \tag{5}$$

where we replaced $N_2$ with $\mathbf{0}$. Importantly, in this approximation the dynamics of species 1 are uncoupled for the dynamics of species 2 and the dynamics of species 2 are purely driven by the periodicity in the matrix entries and the dynamics of species 1.

In this approximation, the resident dynamics (species 1) approaches the periodic solution $N_1^*(t)$. Hence, to understand the fate of the invader (species 2), we only need to consider the periodic matrix model

$$N_2(t+1) = A_2(t, N_1^*(t), \mathbf{0})N_2(t). \tag{6}$$

The long-term population growth rate of species 2 (as an invader) is determined by the dominant eigenvalue $\lambda_{21}$ of the matrix product $\prod_{t=1}^{T} A_2(t, N_1^*(t), \mathbf{0})$. The subscript $_{21}$ denotes the rates of species 2 at the periodic attractor of species 1. Specifically, its invasion growth rate is $r_{21} = \frac{1}{T}\log\lambda_{21}$. If $r_{21} > 0$, then species 2 tends to increase when rare. If $r_{21} < 0$, then species 2 tends to decline exponentially quickly when rare. Similarly, by switching the role of invader and resident, we can define $r_{12} = \frac{1}{T}\log\lambda_{12}$ where $\lambda_{12}$ is the dominant eigenvalue of the matrix $\prod_{t=1}^{T} A_1(t, \mathbf{0}, N_2^*(t))$.

From the mathematical theory of permanence (Roth et al., 2017; Benaïm and Schreiber, 2019), we have the following classification. If $r_{12} > 0$ and $r_{21} > 0$, then the two species coexist in the sense of permanence. If $r_{12} < 0$ and $r_{21} > 0$, then the periodic solution $(\mathbf{0}, N_2^*(t))$ is an attractor and the periodic solution $(N_1^*(t), \mathbf{0})$ is a repellor. This suggests that species 2 could exclude species 1. If the signs of the $r_{ij}$ are reversed, we get the opposite prediction. If $r_{12} < 0$ and $r_{21} < 0$, then both single species periodic solutions are attractors. Hence, there is frequency-dependent effect – which species establishes itself first excludes the other. We note that the exclusion and frequency-dependent effect outcomes are only statements about the local dynamics.

Now, how to compute $r_{21}$? The simplest and most efficient approach is to simulate a modified version of the semi-linear model (5). This modification is a variant of the power method for computing dominant eigenvalues of a non-negative matrix. Let $n_2(t) = \sum_j N_{2,j}(t)$ represent the total population density, $X_2(t) = N_2(t)/n_2(t)$ be the vector of population frequencies, and $\lambda_{12}(t) = n_2(t)/n_2(t-1)$ as the relative change in total population density. In this coordinate system, the semi-linear model becomes

$$\begin{aligned} N_1(t+1) =& A_1(t, N_1(t), \mathbf{0})N_1(t) \\ X_2(t+1) =& \frac{A_2(t, N_1(t), \mathbf{0})X_2(t)}{\|A_2(t, N_1(t), \mathbf{0})X_2(t)\|_1} \\ \lambda_{12}(t+1) =& \|A_2(t, N_1(t), \mathbf{0})X_2(t)\|_1 \end{aligned} \tag{7}$$

where we define $\|v\|_1 = \sum_i |v_i|$ for a vector $v = (v_1, \ldots, v_k)$.

For the initial conditions of this simulation, one can use any non-tribal vector for $N_1(0)$, any probability vector for $X_2(0)$ (a uniform distribution is recommended), and any positive number (1 is recommended) for $\lambda_{12}(0)$. For this modified model,

$$r_{21} = \lim_{t\to\infty} \frac{1}{t} \sum_{s=1}^{t} \log \lambda_{12}(s)$$

Hence, for a sufficiently long simulation of (7), we get approximations of the invasion growth rates by taking the mean of the simulated $\log \lambda_{12}(s)$ values. To determine visually whether a simulation run was sufficiently long, one can plot $\frac{1}{t}\sum_{s=1}^{t} \log \lambda_{12}(s)$ as a function of $t$ and see if it has stabilized or not.

We validated the above method by simulating population dynamics using a small invading population, rather than proportions, until they reach a stable stage distribution, and calculated the invasion growth as the difference between the final and the invading population. These simulations of population dynamics yielded identical results as the numerical methods outlined above.

#### Appendix II. Initial Stage Differences Larger than Generation Time ($|\Delta s| \geq 5$)

In our model in the main text, we did not allow for $|\Delta s| \geq 5$ because we assumed that both species have life histories/phenologies similar enough that they arrive within one generation, and their juvenile stages overlap. However, in some fast-generating systems such as nectar microbiome, the late species may arrive after the early species has reproduced for several generations. Here, we first discuss potential consequences of this scenario, then provide additional simulation results.

##### Potential Consequences of $|\Delta s| \geq 5$

As $|\Delta s| > 5$, one species arrives increasingly late than the other. For simplicity, let's consider a scenario when species 1 arrives early by $\Delta s$ time steps. By definition, $\Delta s > 0$. When species 2 arrives increasingly late, $\Delta s$ could move from the following ranges:

**Range I.** Species 2 arrives before species 1 can reproduce; $\Delta s < 5$.
**Range II.** Species 2 arrives after species 1 reproduces for several generations : $5 \leq \Delta s < t_{eq}$, where $t_e q$ is the time when species 1 reaches a *stable stage distribution*.
**Range III.** Species 2 arrives after species 1 reaches a stable stage distribution: $\Delta s \geq t_{eq}$.

Note that $t_{eq}$ depends on species 1's intraspecific competition. With values we used for simulations in the main text (see Table 1), $t_{eq}$ is likely >> 50, but the exact value also depends on the cutoff threshold at which the stage distribution is considered "stable".

In **Range I**, species 1 has not reproduced yet and is in a stage distribution with only a few juvenile stages, depending on the exact value of $\Delta s$. For example, when species 2 arrives late by two time steps ($\Delta s = 2$), species 1 should only be in $J_1$ and $J_2$ stages. In **Range II**, by the time species 2 arrives, species 1 would have already been in a stage distribution with all stages ($J_1$ to $J_5$, and adults). In this range, species 1 has yet reached a stable stage distribution, meaning that its stage distribution could still change as $\Delta s$ increases, and therefore its interaction with species 2 still changes. In **Range III**, species 1 is at a *fixed* stable stage distribution when species 2 arrives in $J_1$, which is also fixed. Because the two interacting species' stage distributions will not change as $\Delta s$ increases further, their initial interactions will remain the same. However, as $\Delta s$ increases, the time period in which the late arriver is exempt from competition from the early arriver also increases (because the early arriver would have finished early and enter dormancy). If $\Delta s$ is long enough, the temporal overlap between the early and the late arriver would be small, and the two species will likely coexist because neither would receive strong competition from the other.

#### Simulation Results

To test our reasoning, we simulated the scenario with and without stage-mediated interspecific competition, in which $\Delta s$ can reach beyond one generation time. In addition to $2, 4, 8, 16$, and 32 generations per season (as in the main text), we added $T = 64$. For each season length, we simulated the community when $\Delta s/T = 0, \pm 1/12, \pm 1/6, \pm 1/4$, and $\pm 1/3$ . For instance, when $T = 8$ generations, or 48 time steps, $\Delta s$ would be $0, \pm 4, \pm 8, \pm 12, \pm 16$.

Indeed, as seasons get longer, the same proportion of $\Delta s$ gets closer to **Range III**, after which competitive outcomes remain constant. With stage-mediated interspecific competition, when $\Delta s$ is sufficiently large, the two species always coexist (Figure S1A). This is because the late arriver will always have $\Delta s$ time steps during which the early arriver has finished development and enters dormancy, and the longer this time period without competitor, the larger the advantage is for the late arriver. This late arriver advantage is present in the scenario without stage-mediated priority effects but only when season length is extremely long, meaning that a larger $\Delta s$ is necessary for coexistence.

However, the threshold of $\Delta s$ for coexistence appears to be much smaller when $T = 2$ generations: species 1 had an early arriver advantage when it arrived moderately early ($-2 \leq \Delta s \leq 0$) but not when extremely early ($\Delta s = -3, -4$), where it coexisted with species 2 (Figure 3A). Examining the population dynamics of the model, we found that when the early arriver reproduces extremely early ($\Delta s = -3, -4$), the majority of its population is in stage 1 or 2 juveniles; at the same time, all individuals of the late arriving species are in later juvenile stages (4 or 5). During this period, the late arriver has a higher competitive effect on the early arriver due to stage-mediated interspecific competition. However, this period can only last for one to two time steps, and is therefore always shorter than the initial stage difference (three or four time steps). As seasons get longer, both species reproduce to form overlapping cohorts with all stages, and this transient period of late arriver advantage becomes less important.

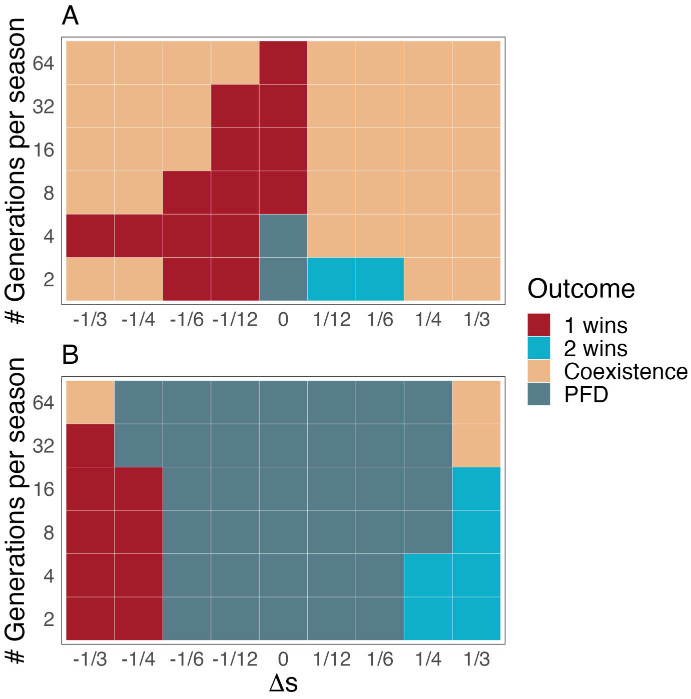


Figure S1. Phase diagrams of competition outcomes across different season lengths ($T$; in number of generations per seasons) and proportion of stage differences to season lengths ($\Delta s/T$). PFD stands for positive frequency dependence (frequency-dependent priority effects). A, with stage-mediated interspecific competition; B, without stage-mediated interspecific competition. See Table 1 for other parameters used.

### Appendix III. Supplemental Figures

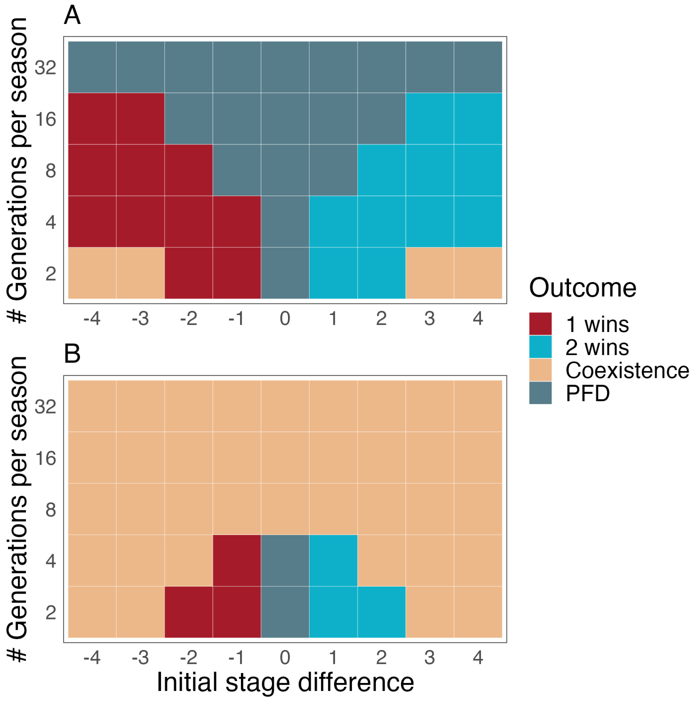


Figure S2. Phase diagrams of competition outcomes across different season lengths ($T$; in number of generations per seasons) and stage differences ($\Delta s$). Parameters used: A, $\alpha_{11} = \alpha_{22} = 0.05$; B, $\alpha_{11} = \alpha_{22} = 0.06$. PFD stands for positive frequency dependence (frequency-dependent priority effects). See Table 1 for other parameters used.

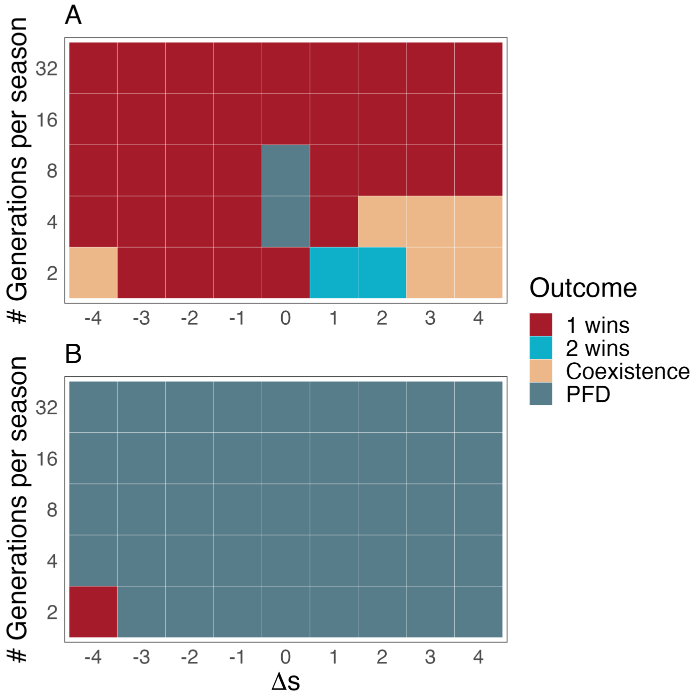


Figure S3. Phase diagrams of competition outcomes across different season lengths ($T$; in number of generations per seasons) and stage differences ($\Delta s$) when adults are allowed to compete with juveniles, and adult mortality and fecundity are also density-dependent. Parameters used: A, $\alpha_{11} = \alpha_{22} = 0.05$; B, $\alpha_{11} = \alpha_{22} = 0.06$. PFD stands for positive frequency dependence (frequency-dependent priority effects). See Table 1 for other parameters used.

### Appendix IV. Stage-Mediated Intraspecific Competition

Multiple studies have shown that the ecology of conspecific individuals at different ontogenetic stages can be substantially different, creating the potential of stage-mediated intraspecific competition (Miller and Rudolf 2011; Rudolf and Rasmussen 2013a,b; Rudolf et al. 2014; Murillo-Rincón et al. 2017). Therefore, we incorporated stage-mediated intraspecific competition in addition to stage-mediated interspecific competition (Eqn. 1 in Methods) and compared results with those presented in the main text. Briefly, the qualitative difference between scenarios with and without stage-mediated competition persisted, and adding stage-mediated intraspecific competition lowered the overall strengths of intraspecific competition for both species, and led to general shifts in long-term competitive outcomes.

#### Deriving Stage-Mediated Intraspecific Competition Coefficients

Similar to Eqn. 1 in the main text, intraspecific competition between stage $i$ and $j$ of species 1 is calculated by:

$$\alpha_{1,i|1,j} = B_{11}/(1 + \exp(d - (j - i)/c) \tag{S1}$$

All else being equal, $B_{11}$ denotes a different maximum competition coefficient from $B$, which is the maximum interspecific competition coefficient. Here, individuals of earlier stages experience a higher intraspecific competition from later stages, and this applies to both species (Figure S4). We explored a range of $B_{11}$ and $B_{22}$ by multiplying intraspecific competition coefficients used in constant (non-stage-mediated) cases by two; this ensured that for individuals of the same stage, their intraspecific competition coefficient is $B_{11}/2$ or $B_{22}/2$, the same coefficient used in constant cases.

#### Results

Comparing with scenarios with equivalent intraspecific competition coefficients in the main text and in Appendix III, adding stage-mediated intraspecific competition strongly promoted positive frequency dependence, suggesting that the overall intraspecific competition was much smaller comparing to interspecific competition in the system. For example, when $\alpha_{11} = \alpha_{22} = 0.06$ without stage-mediated intraspecific competition, overall competitive outcomes trended strongly towards coexistence interspecific competition was generally smaller than intraspecific competition (Figure S3). In the equivalent scenario where the maximum intraspecific competition coefficients

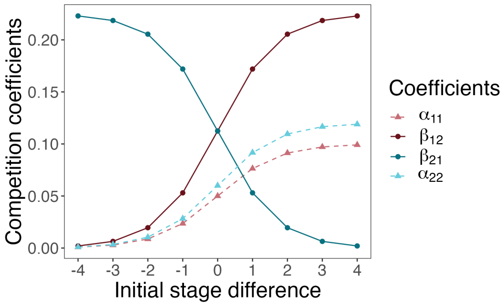


Figure S4. The scaling function of stage-mediated intra- and interspecific competition. All four per-capita competition coefficients are determined by $\Delta s$. The function assumes an early arriver advantage. Parameters used: for intraspecific competition, $B_{11} = 0.10$, $B_{22} = 0.12$; for interspecific competition, $B = 0.225$; in both cases $c = 0.85$, $d = 0$.

$B_{11} = B_{22} = 0.12$, the numeric average of the coefficients was the same as $\alpha_{11}$ and $\alpha_{22}$ (0.06; see Figure S4), but the overall competitive outcomes trended strongly towards positive frequency dependence, indicating smaller intraspecific than interspecific competition. Allowing for stage-mediated intraspecific competition decreased the intraspecific competition (and therefore overall competition) experienced by larger stages, which are closer to reproductive adults, and this decrease likely promoted fecundity, subsequently decreasing the overall intraspecific competition. In addition, because our calculation of invasion growth ignores the intraspecific competition of the invader (see **Methods**), changes in competition outcomes were only caused by the resident's equilibrium population and stage structure.

To further explore the effect of stage-mediated intraspecific competition, we adjusted both the maximum intraspecific competition coefficient $B_{11}$ and $B_{12}$, and the scaling factor $c$ in Eqn. S1 to examine whether we could recreate similar phase diagrams as in the main text, where the model only included stage-mediated interspecific competition. Increasing $B_{11}$ and $B_{22}$ increased the overall intraspecific competition, and increasing $c$ led to less variations of intraspecific competition coefficients with interacting stages (Figure S6). With this set of parameters, patterns of long-term competitive outcomes resembled those in the model with only stage-mediated interspecific competition (Figure 3A).

Together, we concluded that stage-mediated intraspecific competition did not have an interactive effect with stage-mediated interspecific competition, but only an additive effect; it neither contributed nor diminished stage-mediated priority effects observed in the model.

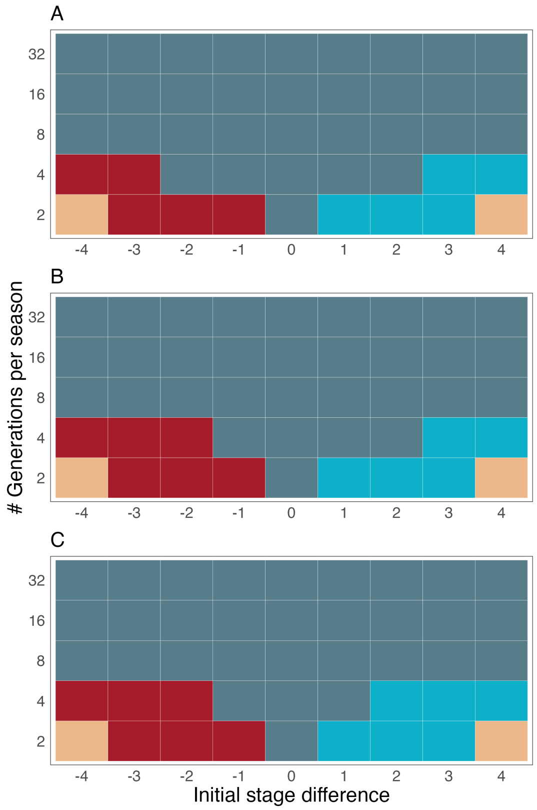


Figure S5. Phase diagrams of competition outcomes across different season lengths ($T$; in number of generations per seasons) and initial stage differences ($\Delta s$), with both stage-mediated intra- and interspecific competition coefficients. Each panel uses a different set of intraspecific competition coefficients: A, $B_{11} = B_{22} = 0.10$, which is equivalent to $\alpha_{11} = \alpha_{22} = 0.05$ in Appendix III (Figure S2); B, $B_{11} = 0.10, B_{22} = 0.12$, which is equivalent to $\alpha_{11} = 0.05$, $\alpha_{22} = 0.06$ in the main text; C, $B_{11} = B_{22} = 0.12$, which is equivalent to $\alpha_{11} = \alpha_{22} = 0.06$ in Appendix III (Figure S2). PFD stands for positive frequency dependence (frequency-dependent priority effects). See Table 1 for other parameters used.

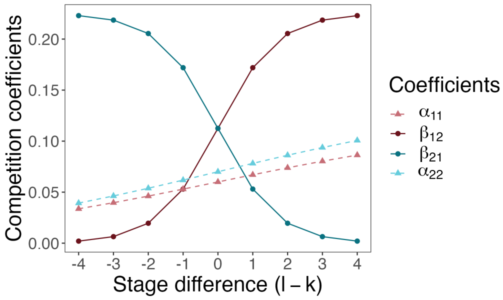


Figure S6. The scaling function of stage-mediated intra- and interspecific competition. All four per-capita competition coefficients are determined by $\Delta s$. The function assumes an early arriver advantage. Parameters used: for intraspecific competition, $B_{11} = 0.12$, $B_{22} = 0.14$, $c = 4.25$, $d = 0$.

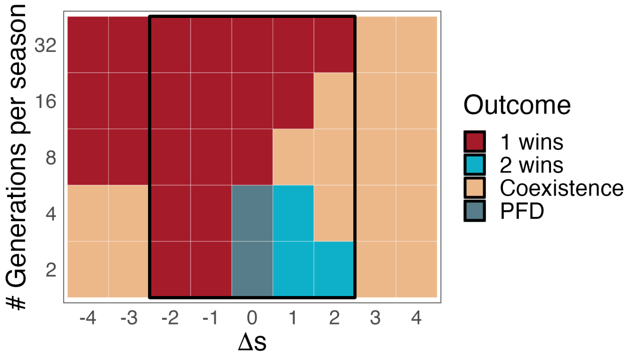


Figure S7. Phase diagrams of competition outcomes across different season lengths ($T$; in number of generations per seasons) and initial stage differences ($\Delta s$), with both stage-mediated intra- and interspecific competition coefficients and adjusted parameters. In all cases, $B_{11} = 0.12$, $B_{22} = 0.14$, and $c = 4.25$. PFD stands for positive frequency dependence (frequency-dependent priority effects). See Table 1 for other parameters used.
